# The Gp78 ubiquitin E3 ligase drives mitophagy by regulating omegasome formation

**DOI:** 10.64898/2026.09.04.749530

**Authors:** Milene Ortiz-Silva, Bharat Joshi, Jerry Zheng, Ren Takimoto, Sanchit Chopra, Nozomu Yachie, Ghassan Hamarneh, Ivan R. Nabi

## Abstract

Autophagy is initiated at endoplasmic reticulum (ER)-associated omegasomes, yet how the ER contributes to mitochondrial cargo degradation during early autophagosome formation remains unclear. Gp78 is an ER-resident RING E3 ubiquitin ligase involved in ER-associated degradation (ERAD) that also promotes Parkin-independent mitophagy. However, the spatial relationship between Gp78 ubiquitin ligase activity, omegasome formation, and mitochondrial degradation has yet to be defined. Combining targeted mitochondria labeling and super-resolution STED microscopy, we show that the Gp78 ubiquitin ligase is localized to and regulates DFCP1-positive omegasome formation, where it drives mitochondrial protein breakdown. Using fluorescent mitophagy reporters and acute ivermectin-induced mitophagy, generation of mitophagy intermediates lacking mitochondrial outer membrane (TOMM20) was found to be dependent on the Gp78 RING finger as well as the BAG6–UBL4A–USP13 ubiquitin-ligase module, and independent of PINK1 and Parkin. Object-based analysis of 3D STED images further showed that both DFCP1- and LC3-positive structures associate primarily with intact mitochondrial fragments which show progressive degradation of mitochondrial content. Gp78 localizes to and promotes, via BAG6, the formation of DFCP1-positive omegasomes and mitochondrial protein degradation in these early autophagic structures. ER-derived omegasome biogenesis is therefore coupled to Gp78-dependent ubiquitin ligase activity and local degradation of mitochondrial cargo during mitophagy.

## Introduction

Macroautophagy/Autophagy is a process by which the cell targets its own components for degradation. Autophagy can be induced upon cellular starvation or other cellular stresses, providing nutrients for cell survival. Selective autophagy targets damaged cellular organelles for disposal, such as mitochondria via mitophagy [1]. Mammalian autophagosome (AP) initiation is mediated by recruitment of membrane from a cytoplasmic platform, most commonly the endoplasmic reticulum (ER), to form an isolation membrane or omegasome [2–4] which can be continuous with or form close contacts with the ER [5,6]. Omegasome formation is triggered by the Atg1/Ulk1 kinase complex that recruits class III phosphatidylinositol 3-kinase complex I (PI3KC3-C1), generating phosphatidylinositol 3-phosphate (PtdIns3P)-enriched ER subdomains that recruit the PI3P-binding FYVE domain containing protein DFCP1 [5,7–9]. DFCP1 therefore labels the ring-or cup-shaped omegasome structures from which the PG expands through recruitment of the ubiquitin-like ATG12–ATG5–ATG16L1 complex, which mediates lipidation of MAP1LC3/LC3-I to membrane bound LC3-II, PG expansion and formation of the double-membraned AP [4]. AP fusion with endolysosomes results in the formation of acidic autolysosomes containing degradative lysosomal hydrolases and cargo degradation [10].

The endoplasmic reticulum-associated degradation (ERAD) pathway plays a major role in the quality control of terminally misfolded proteins in the ER by coupling their retrotranslocation to cytosolic ubiquitylation, and subsequent proteasomal degradation [11]. Gp78 (also known as autocrine motility factor receptor - AMFR) is an ER-resident RING E3 ubiquitin ligase and a central organizer of one branch of the mammalian ERAD pathway [12–17]. Gp78 activity is fine-tuned through reversible regulation of the cytosolic BAG6 chaperone complex. Specifically, Gp78-mediated ubiquitylation of the BAG6 complex subunit UBL4A transiently impairs BAG6 function, whereas the Gp78-associated deubiquitinase USP13 reveres this modification, thereby maintaining BAG6 activity and efficient ERAD [17]. Gp78 ubiquitin ligase activity has also been implicated in PRKN/Parkin-independent mitophagy [18–21]. Consistent with this, wild-type Gp78, but not a RING finger-defective mutant, promotes mitochondrial fragmentation by mediating the ubiquitylation and proteasome-dependent degradation of mitofusins, with MFN1 identified as a key substrate required for Gp78-induced mitophagy [19,21]. Knockout of Gp78 in HT-1080 fibrosarcoma cells impaired basal mitophagy by preventing the recruitment of LC3-positive autophagosomes to depolarized mitochondria, resulting in compromised mitochondrial health and elevated mitochondrial ROS [18].

Mookherjee and colleagues [20] proposed a coupled “reticulo-mitophagy” mechanism in which Gp78-dependent degradation of outer mitochondrial membrane (OMM) proteins allows the ER-phagy receptor RETREG1/FAM134B to engage OPA1 and promote autophagic turnover of Gp78, OPA1 and other mitochondrial components. Consistently, Gp78 was shown to ubiquitinate FAM134B to promote its clustering, LC3B engagement and ER-phagy flux [22]. Here, combining 2D and 3D super-resolution STED microscopy of fluorescent mitophagy reporters and acute ivermectin-induced mitophagy, we localize Gp78 to DFCP1-positive omegasomes where it drives mitochondrial protein breakdown and is required for mitophagy.

## Results

### Ivermectin-induced mitophagy is regulated by the Gp78/USP13/BAG6 machinery

Gp78 knockout HT-1080 cells have deficient basal mitophagy [18]. To study acute induction of mitophagy, we utilized Ivermectin (IVM), a macrocyclic lactone and antiparasitic agent identified as a potent inducer of mitophagy [23]. Unlike classical ionophores (CCCP) or oxphos inhibitors (oligomycin/antimycin), that induce mitophagy through mitochondrial depolarization, IVM induces mitochondrial stress through fission-induced fragmentation, making it a unique tool to study early mitophagy kinetics. Treatment of HT-1080 Gp78 WT cells with 10µM IVM led to a robust and time-dependent induction of the autophagic marker LC3-II (Fig. 1A) that was not observed in Gp78 KO cells, consistent with previous findings that HT-1080 cells rely on Gp78 for both basal and induced mitophagy [18]. To monitor mitophagy, we expressed a mitochondria targeted GFP attached to the Cox4 matrix targeting sequence, called mito-GFP (Fig. 1B), and labeled the OMM with anti-TOMM20 antibody. High-resolution 2D-STED imaging revealed that IVM induced extensive mitochondrial fragmentation and the emergence of mitochondrial structures that retained matrix-GFP signal but showed partial or complete loss of the OMM marker (Fig. 1B). We termed these structures “mitoAVs” (mitochondrial autophagic vesicles), representing pre-lysosomal mitophagy intermediates where the OMM has been lost, but that retain matrix localized mitoGFP that has yet to be quenched by lysosomal acidity.

**Fig. 1.**
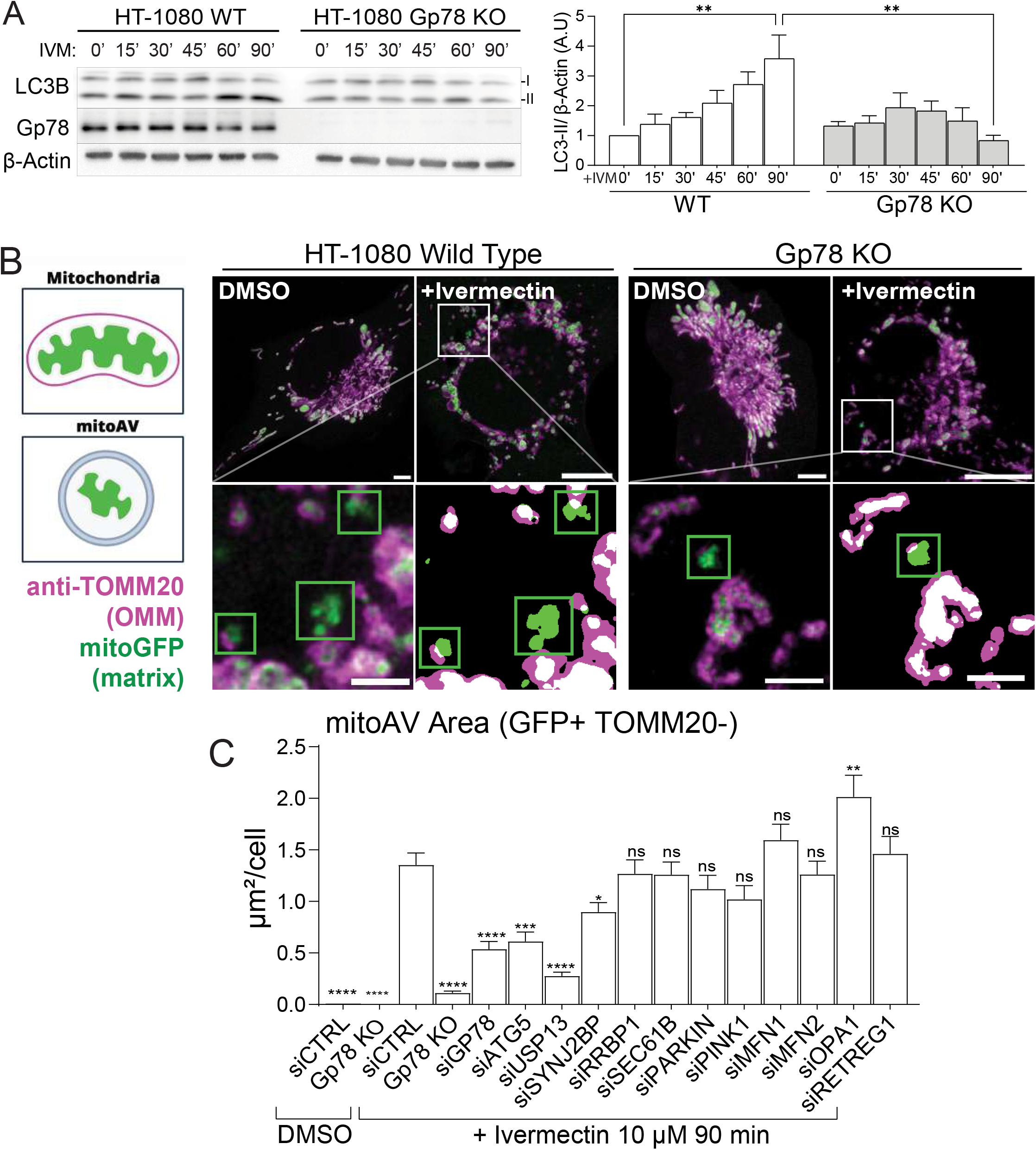
Gp78 is required for acute ivermectin-induced mitophagy. A. WT and Gp78-KO HT-1080 cells were treated with 10 µM ivermectin (IVM) for 0–90 min and immunoblotted for LC3B, Gp78 and β-actin. LC3-II abundance normalized by β-Actin quantified at right. B. Representative scheme of mitoAVs identification and 2D-STED images of cells expressing matrix-targeted mitoGFP (green) and immunolabeled for the OMM protein TOMM20 (magenta). Insets of identified mitoAVs and corresponding segmentation masks are shown below. Scale bars, overviews 5 µm; insets 1 µm. C, Total MitoAV area per cell in DMSO-treated siCTRL and Gp78-KO cells and in IVM-treated siCTRL, Gp78-KO, siGp78, siATG5, siUSP13, siSYNJ2BP, siRRBP1, siSEC61B, siPARKIN, siPINK1, siMFN1, siMFN2, siOPA1 and siRETREG1 conditions. Data are mean ± SEM from n ≈ 50 cells across 3 independent experiments. Statistical comparisons were made against wild type siCTRL + IVM90 using Anova one-way and Tukey post hoc. ns, not significant; *P < 0.05, **P < 0.01, ***P < 0.001 and ****P < 0.0001.

To map the molecular machinery required for mitoAV formation, we performed a targeted siRNA mini-screen and quantified the total mitoAV area per cell using an automated thresholding and masking pipeline (Fig. 1C). Consistent with our visual observations, Gp78 knockdown or knockout resulted in a significant reduction in mitoAV area upon IVM treatment, confirming Gp78 as a primary regulator of mitophagy in HT-1080 cells [18]. Similarly, knockdown of the core autophagy factor Atg5 significantly inhibited mitoAV accumulation, indicating that these structures are dependent on the canonical autophagic machinery. Knockdown of the mitophagy factors PINK1 or PRKN/Parkin had no significant effect on IVM-induced mitoAV area, consistent with the Parkin-independence of Gp78-dependent mitophagy [19]. Gp78 also promotes the formation of ribosome-studded mitochondria-ER contacts or riboMERCs [24,25]. However, knockdown of riboMERC tethers RRBP1-SYNJ2BP [26] showed that loss of ER-localized RRBP1 had no effect on mitophagy while knockdown of the mitochondrial SYNJ2BP partner had a significant but limited effect on mitophagy. This suggests that riboMERC formation is not critically required for Gp78 mitophagy function OPA1 depletion increased mitoAV area; this may reflect reduced mitochondrial fusion amplifying IVM-induced fragmentation and increasing the amount of mitochondrial material entering mitoAVs. Of particular interest, knockdown of the de-ubiquitinase USP13 dramatically limited mitoAV formation, indicating of a role for the Gp78 ubiquitin ligase ERAD module in Gp78-dependent mitophagy [17].

Using this approach, we were unable to detect basal mitoAV formation or Gp78-dependent basal mitophagy in HT-1080 cells, which led us to use the more sensitive SU9-mGFP-HaloTag mitophagy probe [27] (Fig. 2A). The SU9 protein is an IMM resident protein, and when expressed by the construct is imported to the mitochondria matrix. The construct carries both a GFP, quickly degraded after delivery to the lysosome, and a HaloTag, that when bound to its ligand is more resistant and accumulates as a detectable processed fragment. This allows SU9-mGFP-HaloTag to capture downstream mitophagic events not observable due to mito-GFP quenching in the acidic lysosomal environment. Paired with TOMM20 antibody labeling, healthy mitochondria are positive for TOMM20, SU9-GFP and SU9-Halo. MitoAVs are identified as SU9-GFP positive structures partially or completely devoid of TOMM20, whereas late-stage mito-lysosomes, henceforth called “mitoLYs”, are characterized as HaloTag-positive and GFP-negative structures, often also losing TOMM20. Using the SU9-GFP-Halo probe with TOMM20 labeling, both mitoAVs and mitoLYs were detected under basal (DMSO) and damage-induced (IVM 90 min) conditions. Upon IVM treatment, there is a clear increase in mitoAV total area, count, and size while mitoLYs show an increase in total area and count, but no significant change in size (Supplementary Fig. 1).

**Fig. 2.**
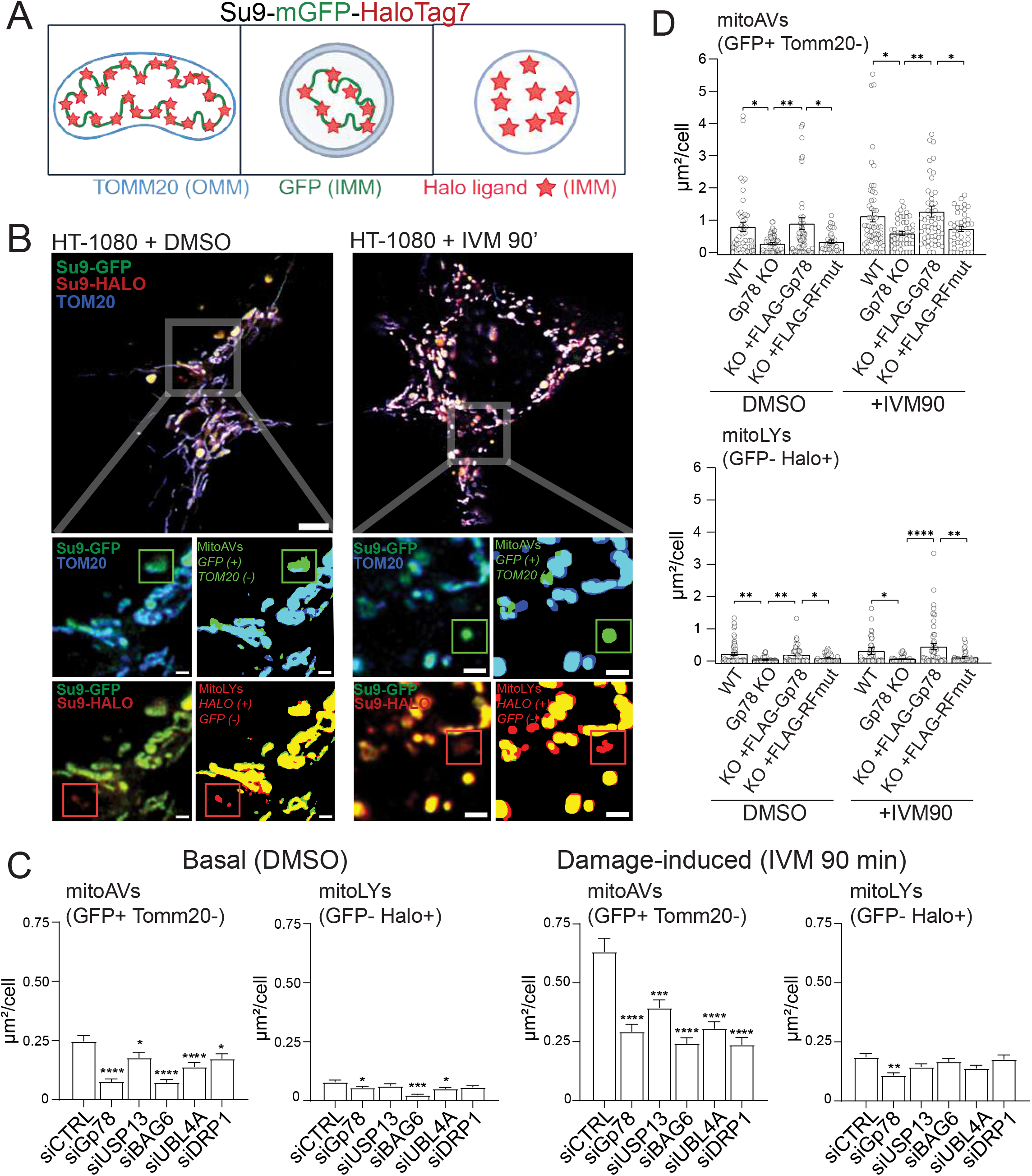
The Gp78 BAG6–UBL4A–USP13 ubiquitin-ligase module regulates basal and ivermectin-induced mitophagy phenotypes. A. Schematic of the dual SU9-HaloTag7-GFP mitochondrial reporter: Healthy mitochondria retain TOMM20, GFP and Halo-ligand signals; MitoAVs retain GFP and Halo signal but have lost or strongly reduced TOMM20; MitoLYs retain the acid-stable Halo-ligand signal but have lost GFP fluorescence. B. Representative 2D-STED images of the SU9 reporter-expressing HT-1080 cells treated with DMSO or 10 µM IVM for 90 min. SU9-GFP is green, SU9-Halo is red, and TOMM20 is blue. Enlarged regions (left) and segmentation masks (right) identify GFP-positive/TOMM20-negative MitoAVs and GFP-negative/Halo-positive MitoLYs. Colors are separated into blue/green and green/red in the inset for ease of view. Scale bars, overviews 5 µm; enlarged images 1 µm. C. Quantification of MitoAV and MitoLY total area per cell after gene silencing with siCTRL, siGp78, siUSP13, siBAG6, siUBL4A or siDRP1 under basal (DMSO) and damage-induced (IVM, 90 min) conditions. DData are mean ± SEM from n ≈ 50 cells from 3 independent experiments. Statistical comparisons were made against siCTRL using Anova one-way and Tukey post-hoc. ns, not significant; *P < 0.05, **P < 0.01, ***P < 0.001 and ****P < 0.0001.

Knockdown of DNM1L/DRP1, which inhibits mitochondrial fission, prevented mitoAV formation (Fig. 2C), consistent with its requirements for IVM-induced [23] and Gp78-dependent mitophagy [18] and with increased mitoAV area observed upon knockdown of the mitochondrial fusion protein OPA1 (Fig. 1C). siGp78 reduces mitoAV formation under basal and damage-induced conditions but mitoLY area and count only upon IVM treatment (Fig. 2C). The more sensitive detection of mitoAVs may be due to the stable accumulation of mitoLYs over time, limiting detection of IVM-induced mitoLYs. Knockdown of BAG6, required for Gp78 activity [28], prevents both mitoAV and mitoLY formation under basal and damage-induced conditions. Knockdown of UBL4A, but not USP13, prevented basal formation of mitoAVs, and both were required for damage-induced formation of mitoAVs and mitoLYs. HT-1080 mitophagy was dependent on Gp78 ubiquitin-ligase activity as rescue of Gp78 KO HT-1080 cells by tet-inducible restoration of wild-type Gp78, but not its Ring-finger mutant (RFmut), restored IVM induction of mitoAVs, and therefore mitophagic activity (Fig. 2D). These data identify SU9-GFP-Halo as a sensitive reporter for basal and damage-induced mitophagy and define a critical role for the BAG6-UBL4A-USP13 complex in Gp78-dependent mitophagy.

### Mitochondrial degradation occurs in the omegasome

To investigate at which stage of the mitophagic process Gp78 activity impacts mitophagy, we transfected SU9-GFP-Halo stable HT-1080 cells with either DFCP1-mCherry to label omegasomes [5] or LC3-RFP to label elongating phagophores and fully formed autophagosomes [29]. Cells were subjected to an IVM time course (0, 30, 60, and 90 min) and then fixed and labeled for the OMM marker TOMM20. 3D STED volumetric images were acquired, and TOMM20/SU9-GFP-Halo labeling was analyzed as described previously to identify mitoAVs and mitoLYs (Fig. 3A). Representative images show how the intact mitochondrial population and the autophagic vacuole population (mitoAV and mitoLY) interact with DFCP1 omegasomes and LC3 labeled phagophores and autophagosomes (Fig. 3A, Supp Videos 1, 2, 5, 6). Both DFCP1 and LC3 populations present structures of varied size and shape. Similar to mitoAVs and mitoLYs, DFCP1-positive structures increase in number significantly over time of IVM treatment while the number of LC3 structures remains constant, the apparent reduction in number of LC3 structures upon IVM 30- and 60-minute treatment is not significant (Fig. 3B). We observed that DFCP1 structures were often found adjacent to mitochondria. To quantify these docking events, we counted the number of DFCP1 and LC3 structures overlapping TOMM20 (OMM) but not SU9-GFP or SU9-Halo (matrix). Docking events for DFCP1, but not LC3, showed a significant increase at 30 minutes post-IVM treatment suggesting that omegasome docking with mitochondria is an early and transient event in the mitophagy process.

**Fig. 3.**
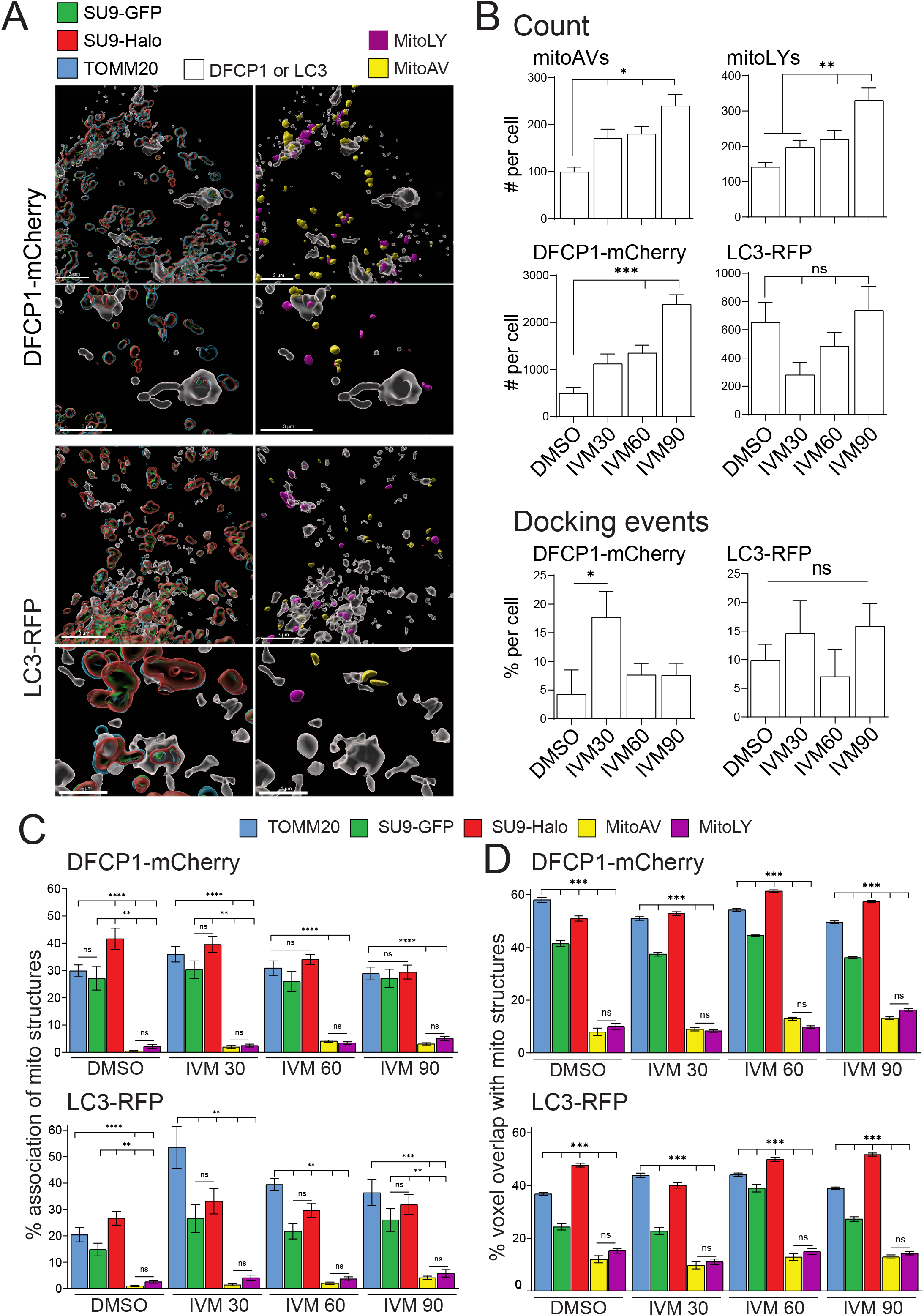
DFCP1-and LC3-positive structures associate predominantly with intact mitochondria fragments. A. Representative 3D-STED surface renderings of SU9 reporter-expressing HT-1080 cells co-expressing DFCP1-mCherry or LC3-RFP and immunolabeled for TOMM20 after IVM 90 minutes. ‘Mitochondria views show TOMM20 (blue), SU9-GFP (green) and SU9-Halo (red) on the left; AV views show segmented masks of MitoAVs (yellow) and MitoLYs (magenta) on the right. For both views, DFCP1 or LC3 structures are shown in transparent white. Rotational views shown in Supplemental Videos 1, 2, 5, 6. B. Number of MitoAVs, MitoLYs, DFCP1-mCherry objects and LC3-RFP objects per cell were quantified during the IVM time course (DMSO, 30, 60 and 90 min). Docking-events represent the number of DFCP1 or LC3 objects per cell overlapping OMM TOMM20 and not SU9-GFP or SU9-Halo. C, The percentage association of mitochondrial objects in each class with at least one overlapping DFCP1 or LC3 voxel. D, For each associating mitochondrial object, the percentage of DFCP1 or LC3 voxels overlapping each mitochondrial mask (voxel overlap) was calculated. Data are mean ± SEM from n ≈15 cells from 2 independent experiments. Statistical comparisons were made against every group against each other using Anova one-way and Tukey post-hoc. ns, not significant; *P < 0.05, **P < 0.01, ***P < 0.001 and ****P < 0.0001.

The majority of DCFP1 and LC3 structures show close association with both OMM (TOMM20) and matrix (SU9-GFP/Halo) markers. However, minimal association with either mitoAVs or mitoLYs was observed (Fig. 3A). Object-based colocalization analysis, measuring the number of the indicated mitochondrial labels presenting pixel overlap with DFCP1 or LC3 structures (Fig. 3C) and, for the overlapping structures, the extent of overlap (voxel/voxel) of interacting structures (Fig. 3D), confirms the minimal association of both mitoAVs and mitoLYs with both DFCP1 and LC3. In contrast, a large proportion of DFCP1 and LC3 structures overlap with TOMM20, SU9-GFP and SU9-Halo, suggesting that these early autophagic structures encompass intact mitochondria or portions thereof. pH neutral or poorly acidic mitoAVs and acidic mitoLYs are therefore degradative autophagosomes that form after LC3 release from the autophagosomal membrane.

To determine whether the mitochondrial fragments present in DFCP1-positive omegasomes or LC3-positive autophagosomes undergo degradation, we assessed the overlap between either TOMM20 or SU9-GFP with the more stable SU9-Halo+Ligand reporter (Fig. 4). In contrast to intact mitochondria outside of DFCP1 omegasomes or LC3 autophagosomes, mitochondrial structures within these structures present a highly varied distribution of the three mitochondrial reporters, representing fragmentation and degradation of the engulfed mitochondrial fragments (Fig. 4A, Supp. Videos 3, 4). To quantify this, we determined the ratio of TOMM20 or SU9-GFP colocalizing with SU9-Halo (TOMM20/Halo or GFP/Halo) for intra-DFCP1 or LC3 mitochondria fragments and present the data as a binned gradient. Intra-DFCP1 SU9-Halo-positive fragments lacking TOMM20 or SU9-GFP are heterogeneously distributed over time, containing a range of structures from high degradation (red bin 0-10%) to intact mitochondria fragments (green bin 90-100%). Analysis of mitochondrial fragments showed that the total number of Halo-positive fragments within DFCP1 omegasomes increased over time of IVM treatment, with significant progressive increases of both highly degraded and intact fragments (Fig. 4B). Proportional analysis, presenting the degradation bins as a proportion of the total number of omegasome-encompassed mitochondrial fragments, showed that there was a significant increase in highly degraded fragments and decrease of intact TOMM20/SU9-Halo fragments, reflecting OMM degradation, but not of SU9-GFP/SU9-Halo structures in omegasomes over time. Mitochondrial fragments within LC3 autophagosomes presented a similar degree of degradation that showed minimal changes over time of IVM treatment (Fig. 4 A,B). The significant change between IVM30 and IVM90 relates to the reduction of fragments detected at IVM30, likely due to the impact of IVM on stable LC3 autophagosomes. These data suggest that there is progressive loss of the OMM marker with IVM treatment within omegasomes.

**Fig. 4.**
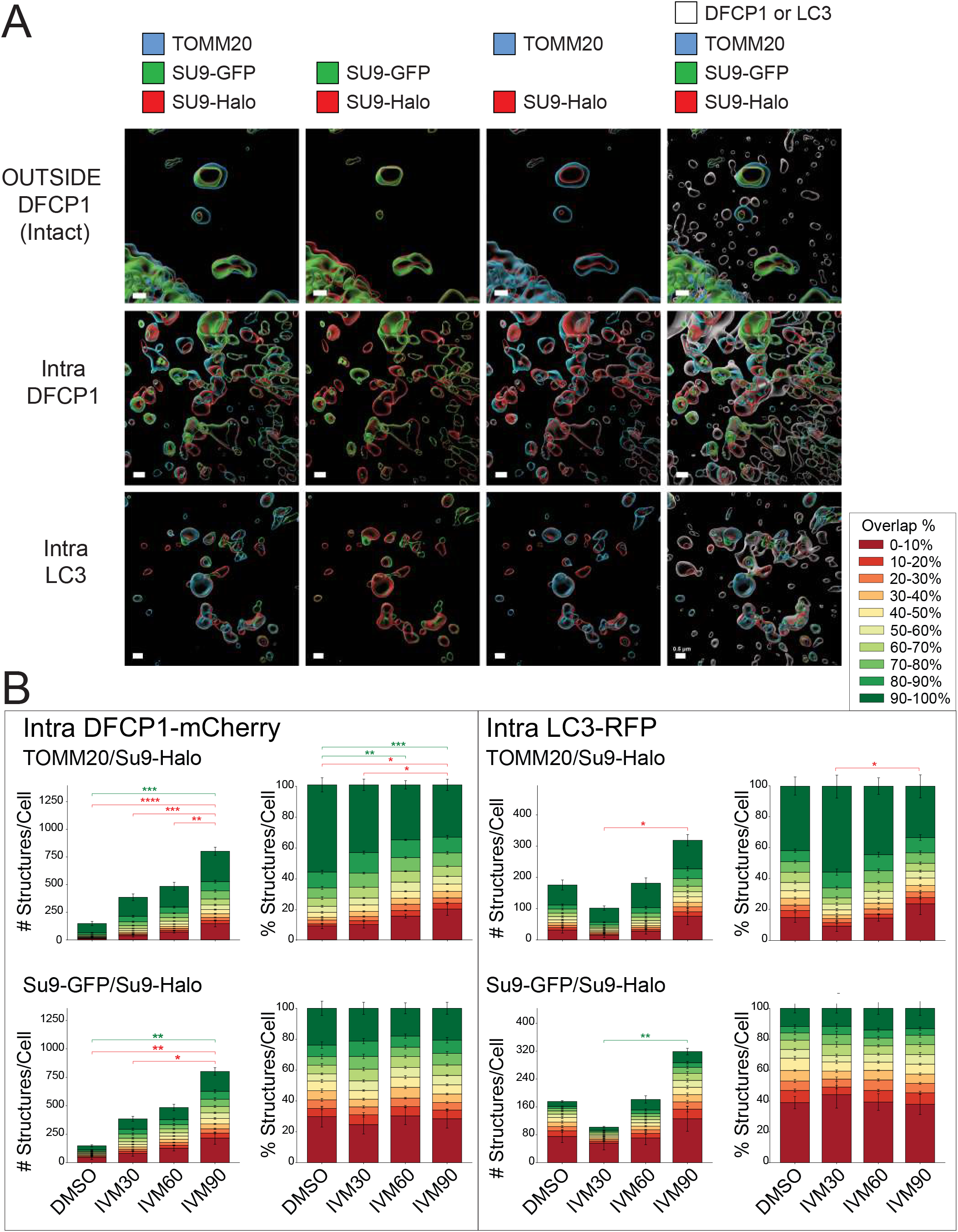
Mitochondrial fragmentation within DFCP1-positive omegasomes and LC3-positive structures. A. Representative 3D-STED surface renderings of mitochondrial objects outside DFCP1 structures and representation of mitochondria channels inside DFCP1-mCherry-or LC3-RFP-positive structures after IVM 90 min. TOMM20 is blue, SU9-GFP is green, SU9-Halo is red and DFCP1 or LC3 is white. Colors are separated into green/red/blue, green/red or blue/green to highlight extent of overlap between the different channels. Merged channel views illustrate heterogeneous retention of the three mitochondrial markers within DFCP1-positive omegasomes and LC3-positive AVs. Scale bar = 0.5 µm. Rotational views shown in Supplemental Videos 3-4. B. SU9-Halo-positive mitochondrial fragments within DFCP1 or LC3 structures were binned in 10% intervals according to the percentage of TOMM20/SU9-Halo or SU9-GFP/SU9-Halo overlap. Stacked bin bars show the number of structures per cell on the left and the proportional distribution on the right, after DMSO or 30, 60 or 90 min of IVM treatment. Dark red represents 0–10% overlap (greatest loss of TOMM20 or SU9-GFP relative to SU9-Halo), whereas dark green represents 90–100% overlap (greatest OMM or matrix intactness). Colored significance brackets refer to the correspondingly colored overlap bin. Data are mean ± SEM from n ≈ 15 cells from 2 independent experiments. Statistical comparisons were made against every group against each other using Anova one-way and Tukey post-hoc; ns = not significant; *P < 0.05, **P < 0.01, ***P < 0.001 and ****P < 0.0001.

### Gp78 associates with degradative omegasomes

We then tested whether the luminal ER reporter KDEL-Halo or Flag-Gp78 associated with DFCP1-mCherry or LC3-RFP in response to IVM treatment. As seen in Fig. 5A-B, 2D STED analysis shows that DFCP1-mCherry positive puncta colocalized with both KDEL-Halo and Flag-Gp78 under both basal and IVM treated conditions. While IVM treatment did not affect the extent of colocalization of KDEL-Halo with DFCP1, Gp78 colocalization with DFCP1 showed a significant increase with IVM treatment reflecting the recruitment of Gp78 to omegasomes in response to induction of mitophagy. In contrast, LC3-RFP puncta show limited to no association with either ER reporter under both conditions.

**Fig. 5.**
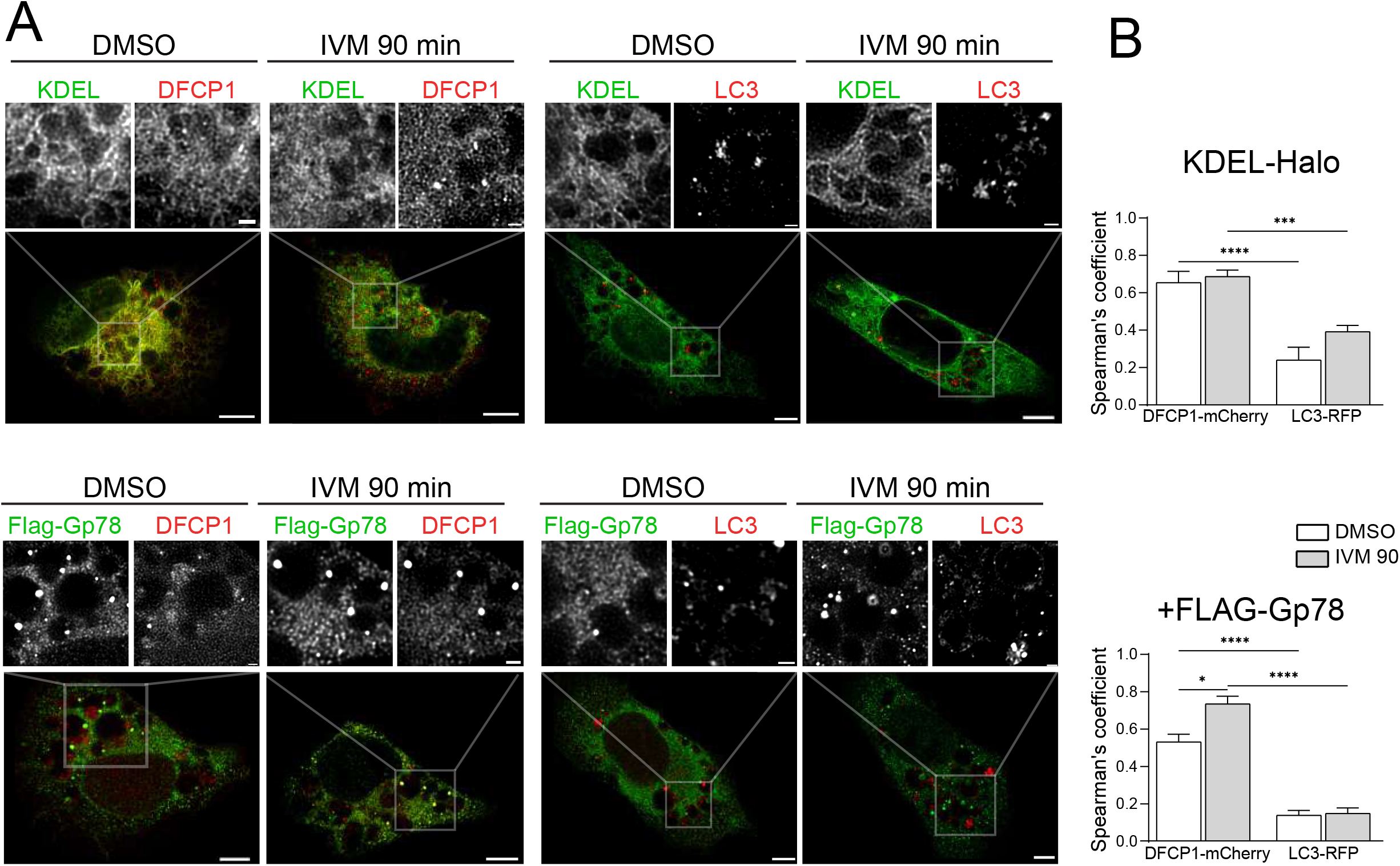
ER and Gp78 preferentially associate with DFCP1-positive structures. A. Representative 2D-STED images of HT-1080 cells co-expressing KDEL-Halo or FLAG-Gp78 (green) with DFCP1-mCherry or LC3-RFP (red), after DMSO or 10 µM IVM treatment for 90 min. Insets show enlarged regions. Scale bars, overviews 5 µm; insets 1 µm. B. Spearman’s correlation coefficients for KDEL-Halo or FLAG-Gp78 with DFCP1 or LC3 under DMSO and IVM conditions. Data are mean ± SEM from n ≈ 36 cells from 3 independent experiments. Statistical comparisons were made against every group against each other using Anova two-way and Tukey post-hoc; relevant significant comparisons are shown. ns, not significant; *P < 0.05, **P < 0.01, ***P < 0.001 and ****P < 0.0001.

To test whether Gp78 can promote mitochondrial degradation in omegasomes, we co-expressed either KDEL-Halo or Gp78-FLAG with DFCP1-mCherry in HT-1080 cells stably expressing the mitochondrial matrix reporter mito-GFP. Cells were treated with IVM for 90 minutes, labeled for TOMM20 and imaged by 3D STED (Fig. 6A, Supp. Videos 7-10). The majority of DFCP1 objects are positive for KDEL or Gp78 and 3D STED views show that DFCP1 omegasomes encompassing mitochondrial fragments are associated with the KDEL-Halo or Flag-Gp78 labeled ER (Fig. 6B). When rendered opaque, the ER channel masks the DFCP1 omegasomes and indicating that omegasomes are localized within the extended ER network.

**Fig. 6.**
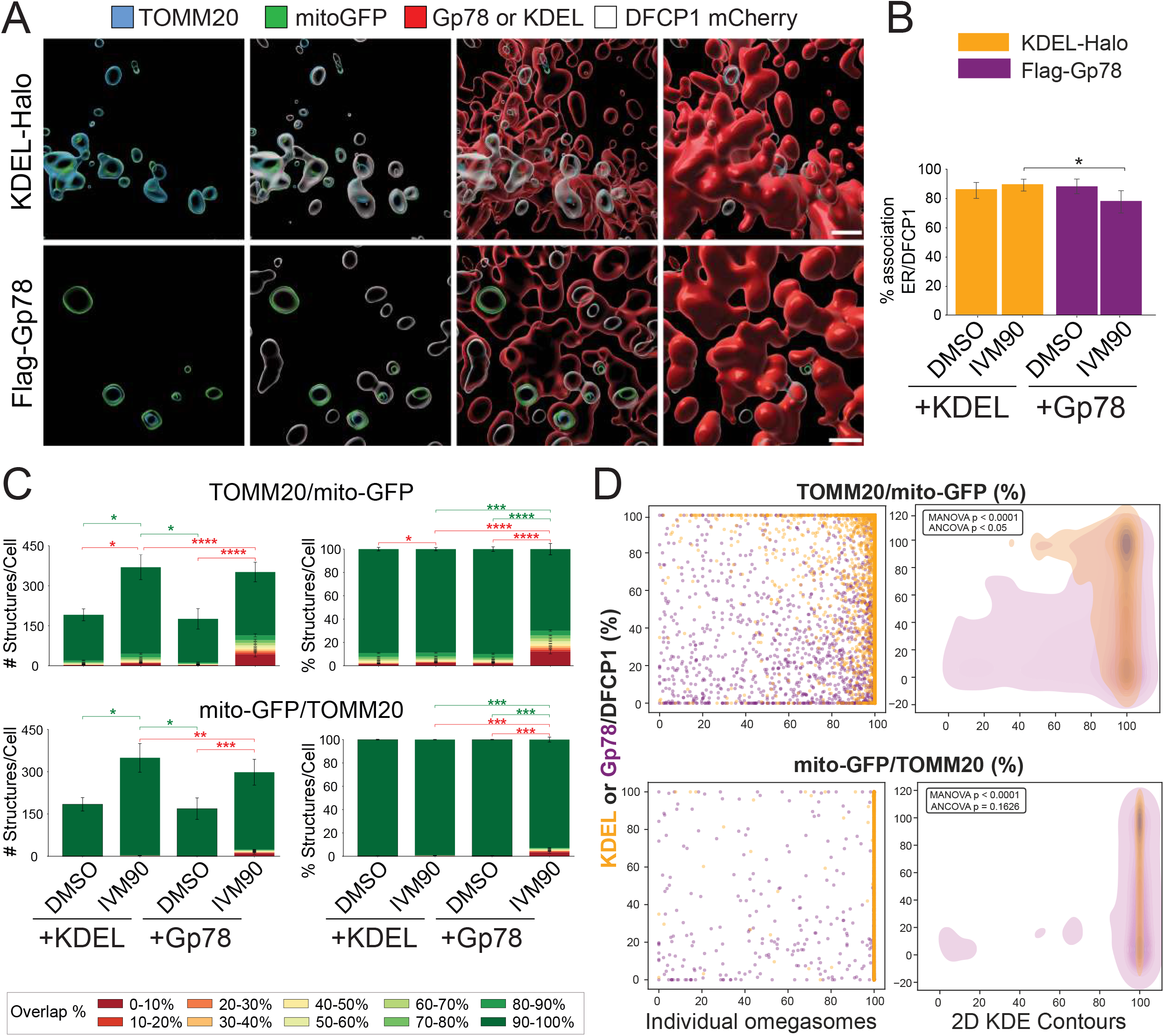
Gp78 abundance induces increased mitochondria degradation inside omegasomes. A. Representative 3D-STED renderings of mitochondrial fragments within DFCP1 omegasomes in HT1080 cells expressing either KDEL-Halo or Gp78 KO cells expressing FLAG-Gp78 after 10 µM ivermectin (IVM) for 90 min. From left to right, the same region is shown with TOMM20 (blue) and mitoGFP (green), with DFCP1-mCherry added (white), with the ER reporter added as a transparent surface (red), and with the ER reporter rendered opaque red, showing embedding of DFCP1 structures within the ER network. Scale Bar = 0.5 µm. Rotational views shown in Supplemental Videos 7-10. B, Percentage of DFCP1 voxels overlapping KDEL-Halo or FLAG-Gp78 in DMSO or IVM90 treated cells. C. DFCP1-associated mitochondrial structures in cells expressing KDEL-Halo or FLAG-Gp78 were binned based on TOMM20/mitoGFP or mitoGFP/TOMM20 percentage overlap after DMSO or 10 µM IVM treatment for 90 min. Stacked bars show the number of structures per cell (left) and the proportional distribution (right) across 10% overlap bins. Colored significance brackets refer to the correspondingly colored overlap bin. Data are mean ± SEM from n ≈ 17 cells from 2 independent experiments. Statistical comparisons were made against every group using Anova two-way and Tukey post-hoc: ns, not significant; *P < 0.05, **P < 0.01, ***P < 0.001 and ****P < 0.0001. D. For each individual intra-DFCP1 mitochondrial fragment in all cells, scatter plots show the KDEL/DFCP1 or Gp78/DFCP1 percent ratio colocalization in the Y-axis and the OMM/mito-GFP (top) or mito-GFP/OMM (bottom) colocalization in the X-axis. These plots present the differential loss of mitochondrial integrity upon overexpression of KDEL-Halo (orange) or Flag-Gp78 (purple). Two-dimensional KDE-contours summarize the population density and centroid position of each marker group. MANOVA was used to test whether KDEL-Halo and Flag-Gp78 structures differed in their joint distribution across the two plotted variables. ANCOVA was used to test whether the relationship between mitochondrial integrity and ER/DFCP1 colocalization differed between marker groups.

We used binning degradation analysis to compare TOMM20/mito-GFP and mito-GFP/TOMM20 percent ratios in omegasomes of cells overexpressing KDEL-Halo or Flag-Gp78. In the absence of the more stable Halo mitochondrial probe, mitochondrial matrix can be completely devoid of GFP signal, therefore we limited the mitoGFP/TOMM20 degradation analysis to the fragments containing both channels, after excluding docking events. Comparison of relative amounts of TOMM20 and matrix mito-GFP provides a measure of TOMM20 and matrix mitochondrial protein degradation in omegasomes. TOMM20 degradation in omegasomes was significantly increased upon IVM treatment in both KDEL and FLAG-Gp78 expressing cells (Fig. 6C). Overexpression of Gp78 enhanced the mitochondrial degradation, as reported by TOMM20/mito-GFP and mito-GFP/TOMM20 ratios, in response to IVM, supporting a role for Gp78 in mitochondrial fragment degradation in omegasomes (Fig. 6C).

We then performed a 2D Kernel Density Estimation (KDE) population analysis to determine the relationship between ER association of DFCP1-labelled omegasomes and mitochondrial degradation percent ratio within these structures being 0% the complete loss of TOMM20 of mitoGFP and 100% complete colocalization, or integral mitochondria. The extent of association (voxel/voxel) of either KDEL-Halo or Flag-GFP with DFPC1 labeled omegasomes was mapped against TOMM20/mito-GFP or mito-GFP/TOMM20 ratios within the same omegasomes in response to IVM (Fig. 6D). Multivariate analysis of variance (MANOVA) revealed a significant 2D spatial divergence between Gp78-positive structures and KDEL-Halo expressing cells for TOMM20/mito-GFP and to a lesser extent for mito-GFP/TOMM20, indicative of Gp78-dependent OMM degradation within omegasomes. Analysis of covariance (ANCOVA, p<0.05) confirmed a significant interaction effect for TOMM20 degradation (i.e. TOMM20/mito-GFP), demonstrating that ER recruitment to DFCP1 omegasomes fundamentally shifts in response to overexpression of Gp78.

To investigate the role of Gp78 ubiquitin ligase activity on omegasome formation and degradation of mitochondrial cargo, we used wild-type and Gp78 KO HT-1080 cells treated with siBAG6 and assessed mitochondrial degradation inside DFCP1-mCherry labeled omegasomes in response to IVM treatment. For both Gp78 KO cells and siBAG6 treated wild-type HT-1080 cells, the number of mitoAvs, mitoLYs and DFCP1-positive omegasomes were dramatically reduced (Fig. 7A). The number of intra-DFCP mitochondria fragments was also reduced, however, we did not observe significant differences in the degradation pattern (0-10% or 90- 100%) within the omegasomes by proportional analysis (Fig. 7B).

**Fig. 7.**
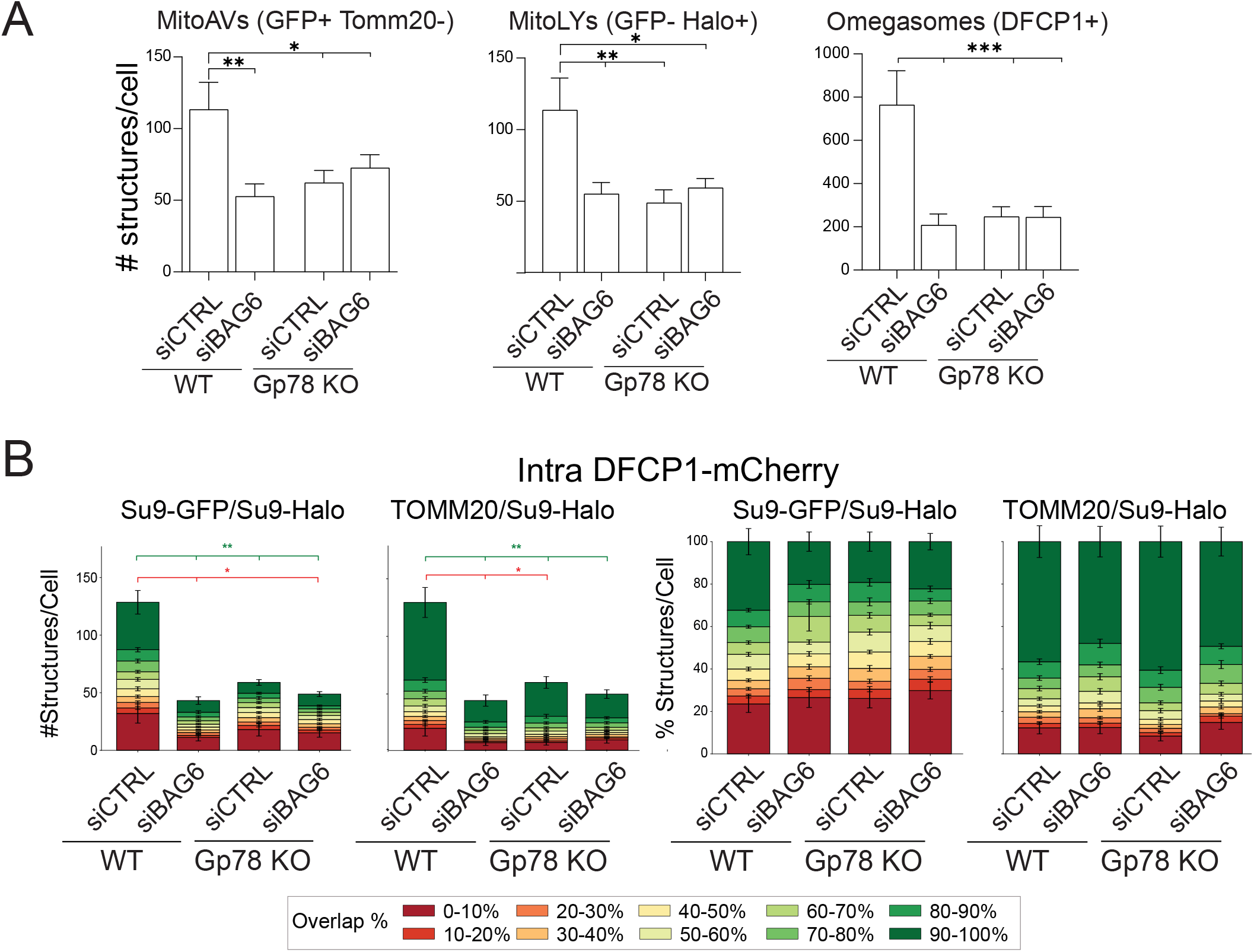
Gp78 and BAG6 promote formation of DFCP1-positive omegasomes. A. Counts of mitoAVs, mitoLYs, and DFCP1-positive structures in HT-1080 WT and Gp78-KO cells treated with siCTRL or siBAG6 after IVM 90 minutes treatment. B, Number and proportional distributions of Halo-positive mitochondrial regions overlapping DFCP1, binned by SU9-GFP/SU9-Halo or TOMM20/SU9-Halo overlap. Colored significance brackets refer to the correspondingly colored overlap bin. Data are mean ± SEM from n ≈ 17 cells across 2 independent experiments. Statistical comparisons were made against every group using Anova two-way and Tukey post-hoc: ns, not significant; *P < 0.05, **P < 0.01, ***P < 0.001 and ****P < 0.0001.

## Discussion

Our findings identify Gp78 ubiquitin ligase activity as an early driver of mitophagy at ER-derived autophagosome initiation sites. The ER is a well-established source of membrane, lipids, and PI3P-enriched domains for omegasome and phagophore formation [5,8,9]. The Gp78 RING-Finger domain that mediates Gp78 ubiquitin ligase activity has been shown to be required for mitophagy induction by Gp78 [18,19,21]. Our data demonstrate that the functional Gp78 ERAD module, composed of BAG6, UBL4A and USP13 [17,30], are required for Gp78-dependent mitophagy and functionally coupled to mitochondrial cargo degradation within omegasomes. Gp78 and BAG6 promote the formation of DFCP1-positive omegasomes and their engulfing of mitochondria fragments. Gp78 ubiquitin ligase activity therefore promotes omegasome formation and thereby mitophagy.

The mitochondrial stressors CCCP and oligomycin/antimycin are commonly used to induce mitophagy, however they require prolonged incubation periods, resulting in heterogeneous induction of mitophagy across the cell population complicating characterization of early intermediates in mitochondrial degradation [31–33]. To resolve early mitophagy events, we applied the rapid, ubiquitin-dependent mitophagy inducer ivermectin (IVM) [23] to HT-1080 cells, which display robust Gp78-dependent basal and damage-induced mitophagy [18]. To monitor mitophagic progression, we used mitochondrially targeted mito-GFP and the more sensitive SU9-GFP-Halo mitophagy probe, that incorporates a SU9-targeted Halotag, that when associated with Halo ligand is highly stable in autolysosomes [27]. Together with OMM labeling for TOMM20, these probes allow us to monitor loss of OMM within autophagic compartments, arrival of mitochondrial cargo to acidic autophagic vacuoles due to quenching of GFP fluorescence (mitoAVs) and identification of mitolysosomes (mitoLYs) identified by the sole presence of SU9-Halo. Consistent with the PINK/Parkin-independence and fission stress mechanism of IVM-induced mitophagy [23], we observed that mitophagy progression was PINK/Parkin-independent and prevented by DRP1 knockdown inhibiting mitochondrial fission. This approach provided an experimental window in which early autophagic events such as ER engagement and omegasome formation and cargo degradation could be examined temporally by 3D super-resolution microscopy during the autophagic maturation process.

The present analysis shows that mitochondria targeted for autophagic degradation enter a Gp78-dependent pathway in which OMM and matrix markers are progressively and asynchronously degraded, before lysosomal acidification. 3D super-resolution imaging of TOMM20 and SU9-GFP-Halo follows cargo degradation through autophagic compartments defined by the autophagic membrane reporters DFCP1 and LC3. Degradative autophagic compartments defined by loss of TOMM20 include GFP-positive mitoAVs and more acidic and degradative, GFP-negative, mitoLYs. DFCP1 and LC3-positive structures associate with intact and partially degraded mitochondrial fragments, but not with mitoAVs or mitoLYs. These results are consistent with omegasomes and phagophores representing biogenetic precursors of more degradative autophagic compartments [4]. ER markers KDEL-Halo and Gp78 colocalize with DFCP1 but not with LC3, confirming a role for ER in omegasome formation and loss during progression to phagophores and early autophagosomes. Our 3D STED analysis supports the formation of DFCP1 enriched domains within the ER that associate with mitochondria upon induction of mitophagy, as evidenced by DFCP1 docking to the OMM. ER-positive omegasomes then engulf mitochondrial fragments and mature to form LC3-positive phagophores and autophagosomes that no longer retain either ER marker. Based on our prior work using tandem-fluorescent LC3 in HT-1080 cells, these LC3-positive intermediates show transient association with low-potential mitochondrial regions [18], presumably before maturing to form degradative autophagic compartments, such as mitoAVs and mitoLYs.

Gp78 ubiquitin ligase activity, through Bag6, is shown here to promote omegasome formation. Gp78-positive ER subdomains associate with DFCP1-positive omegasomes, dock, engulf and promote local degradation of mitochondrial fragments. BAG6 has also been implicated in PINK1-Parkin mitophagy [34] and the general role of this pathway in mitophagy remains to be determined. In Parkin-dependent mitophagy, OMM proteins such as TOMM20, TOMM40, TOMM70, OMP25 and mitofusins undergo ubiquitin-, p97/VCP-and proteasome-dependent degradation before or during autophagic mitochondrial clearance [35–37]. OMM rupture has been proposed to expose IMM components to autophagy machinery and has been shown enable LC3 binding to the IMM receptor prohibitin-2 [38,39]. Here, we detected TOMM20 marker loss inside DFCP1-positive structures and found that Gp78 abundance was associated with greater TOMM20 loss, placing OMM marker loss at the ER-associated omegasome. Gp78 is also implicated in RETREG1/FAM134B mediated ER-phagy [20,22]. RETREG1 was shown to be downstream of Gp78-driven mitochondrial degradation [20]. Consistently, RETREG1 silencing did not inhibit mitophagy in our system suggesting that Gp78 function in mitophagy is distinct from its role in RETREG1/FAM134B-dependent ER-phagy.

Together, our findings describe Gp78 as an early driver of mitophagy within omegasomes. Gp78 interfaces ERAD and mitophagy, with the ER providing the omegasome membrane platform where Gp78 supplies an ubiquitin-dependent degradative activity that generates partially degraded mitochondria for subsequent elimination in degradative autophagic compartments (i.e. mitoAVs and mitoLYs), converts captured mitochondrial cargo into mitoAV and mitoLY intermediates. This positions the omegasome not only as a site of autophagosome biogenesis but also as an early compartment where mitochondrial cargo degradation begins.

## Acknowledgements

This study was supported by research funds from the Canadian Institutes of Health Research (CIHR Project grant AWD-022443). We thank Arun John Peter for helpful discussions.

## Materials and Methods

### Cell lines and culture conditions

Human HT-1080 fibrosarcoma Gp78 wild-type and Gp78 knockout cells were generated by CRISPR-Cas9 as described previously [18]. The Gp78 knockout clone used in this study was G2-41 [18]. Cells were cultured in RPMI supplemented with 10% fetal bovine serum, and 1% L-glutamine (Gibco, Thermo Fisher Scientific). Cells were maintained at 37°C in a humidified atmosphere containing 5% CO and passaged at approximately 30% confluence every 2-3 days using 0.05% trypsin (Gibco, Thermo Fisher Scientific) until passage 7 from initial thawing. Cell lines were authenticated by Short Tandem Repeat (STR) profiling at the TCAG Genetic Analysis Facility (Hospital for Sick Kids, Toronto, ON, Canada www.tcag.ca/facilities/geneticAnalysis.html) and regularly tested for mycoplasma infection by PCR (ABM, Richmond, BC, Canada). When applicable, cells were treated with 10µM of IVM diluted in DMSO for 30, 60 or 90 min, and DMSO was used as vehicle control. When applicable, 1 µM of Live550 Halo ligand (Abberior Instruments GmbH) was pulse-chased 30 minutes before IVM or DMSO treatment and washed with media twice.

### Plasmids, fluorescent reporters and stable cell lines

Mito-meGFP and pMRX-IB-pSU9-HaloTag7-mGFP, mCherry-DFCP1, pmRFP-LC3, Halo-KDEL, were used to label mitochondria matrix, omegasomes, phagophores/autophagosomes, luminal ER, respectively, catalog number can be consulted in Key Resources Table S1. Stable mito-GFP, KDEL-Halo and pSU9-HaloTag7-mGFP HT-1080 lines were generated and selected with geneticin at 500 µg/mL for mitoGFP and/or KDEL-HALO or blasticidin 5 µg/mL for SU9-GFP-Halo for 15 days for selection and later maintained at 10% of each respective concentration. Pooled highly expressing cells were isolated by flow cytometry to establish a stable population.

### DNA and siRNA transfection

Cells were plated at 150,000 cells per 35 mm well approximately 18h before transfection. 400 ng/well of plasmid DNA was introduced using Effectene Transfection Reagen (QIAGEN, Aarhus, Denmark) according to the manufacturer’s instructions. For co-expression experiments, plasmids were combined at a ratio of 1:1. Cells were fixed 24h or 48h after transfection, depending on the experiment. Cells were transfected with siRNAs purchased from Dharmacon/Horizon Discovery; catalogue numbers or sequences are listed in Key Resources Table/Table S1, using Lipofectamine 2000 following the manufacturer’s instructions. Cells were analyzed 48h after siRNA transfection. When DNA and siRNA were used together, cells were co-transfected with 2µg of plasmid DNA and 100pmol siRNA per 35 mm well.

### Rescuing GP78 Knockout with doxy inducible Lentivirus

#### Preparing Plasmids

All primers used for cloning are listed in the accompanying table. The vectors pLV-EF1a-T2A-puro and pLV-dox-Gp78-T2A-lox2272-mTagBFP-lox2272 were constructed using Gibson Assembly. For the pLV-EF1a-T2A-puro vector, SalI- and XhoI digested pLV.CMVenh.gp91.eGFP.cHS4 was combined with: (i) PCR-amplified EF1a promoter (primers RT344/RT345) using pRT193 as the template; (ii) PCR-amplified rtTA-T2A (primers RT340/RT351) using C008_3 as the template; and (iii) PCR-amplified puromycin resistance gene (primers RT352/RT328) using pRT193 as the template. For the construction of pLV-dox-Gp78-T2A-lox66-mTagBFP-lox66, a filler sequence (part of eGFP) was initially inserted at the site of the mutated Gp78, flanked by PaqCI recognition sites with 4-nucleotide overhangs to facilitate subsequent cloning of two different Gp78 variants and the WT Gp78. This intermediate plasmid, designated as pLV-dox-filler-lox2272-mtagBFP-lox2272 was generated by combining SalI-and XhoI-digested pLV.CMVenh.gp91.eGFP.cHS4 with: (i) PCR-amplified tight TRE promoter (primers RT346/478) from pLV.INS-Dox-PaqCI-T2A-eGFP; (ii) PCR-amplified lox2272-BFP-lox2272 (primers RT479/480) from PC302. Three different Gp78 genes were then PCR-amplified separately (primers RT481/482) from plasmids harboring the mutations, each incorporating two PaqCI recognition sites on both edges with overhangs matching the filler plasmid. The PaqCI-digested mutated Gp78 fragments were subsequently ligated into the PaqCI-digested pLV-dox-filler-lox2272-BFP-lox2272 plasmid, resulting in the construction of three plasmids, each containing a Gp78 mutation at a distinct position and WT Gp78. For lentivirus preparation, packaging HEK293T cells were plated either in a 10cm cell culture dish at a density of ∼3×106cells in 10ml of culture medium, 1 day before plasmid transfection. For virus packaging in a 10cm dish, 3.0μg of the transgene vector, 2.25μg of psPAX2 (Addgene, no. 12260), 0.75μg of pMD2.G (Addgene, no. 12259) and 27 μl of 1mgml−1PEI MAX (Polysciences, no. 24765-100) were dissolved in 500μl of Opti-MEM and added to the cell culture. The culture medium was replaced with fresh medium 1day after transfection. Culture medium was collected 24 and 48 h after the initial media change, pooled, and filtered through 0.22μm sterile syringe filters. The lentivirus samples were either concentrated or aliquoted in 500μl volumes into 1.5ml tubes and stored at −80°C. To increase the viral infection titre, collected virus samples were concentrated using a polyethylene glycol (PEG)-based method (https://www.mdanderson.org/research/research-resources/core-facilities/functional-genomics-core/resources.html) with PEG 8000 (Thermo Scientific Chemicals, no. 043443.A3). For concentration with PEG 8000, approximately 20 mL of recombinant virus supernatant was mixed with 6.7 mL of a 4× lentivirus concentrator solution (40% w/v PEG 8000, 1.2 M NaCl, 1× PBS), corresponding to the addition of approximately 2.0 mL of 4M NaCl and 6.7 mL of 1× PBS. The mixture was rotated continuously at 4°C for at least 90 min, then centrifuged at 1,600g for 60min at 4 °C. The supernatant was discarded, and the virus pellet was resuspended in 2 ml of Opti-MEM (Gibco, no. 31985062), achieving a tenfold concentration of the virus sample. The concentrated virus samples were stored at −80 °C.

#### Virus transduction

To introduce rtTA and GP78 variants to cells, parental cells were seeded in 6-well cell culture plates at a density of ∼2 × 105cells per well in 2 ml of culture medium 1 day before transduction. A recombinant virus sample with a 10–100 μL volume of each construct was thawed on ice, mixed with 1.5 μL of 8 μg/ml−1 Polybrene (Sigma-Aldrich, no. TR-1003) and 1.5 ml of fresh culture medium and then applied to the cells. To select transduced cells, the culture medium was replaced with a fresh medium containing 3.0 μg ml−1puromycin (Gibco no. A1113803) 1 day after infection, followed by an additional 3 days of incubation at 1.0 μg ml−1. Cell were then expanded for 2 passages and, when applicable, Gp78 expression was induced with 1 µg/mL doxycycline for 2 hours pulse and 2 hours chase before IVM treatment in Figure 2-D experiments or 16 hours pulse and 2 hours chase in Figure 6 experiments.

#### Cell lysis and immunoblotting

Cells were washed 3 times with ice-cold PBS and lysed in 50 mM Tris-HCl (pH 7.4–7.5), 150 mM NaCl, 1% Triton X-100, and 1 mM EDTA, supplemented with cOmplete™, Mini, EDTA-free Protease Inhibitor Cocktail (Roche/MilliporeSigma). Lysates were incubated for 45 min at 4C and clarified by centrifugation at 10000 rpm x 15 min x 4C. Protein concentration was determined using The Bio-Rad Protein Assay kit (Bio-Rad Laboratories, Hercules, CA, USA). Equal amounts of protein were resolved on SDS–PAGE or gradient gels. Proteins were transferred to PVDF or nitrocellulose as needed semi-dry transfer conditions. Membranes were blocked in 5% milk 1h. Primary antibodies were incubated at 1:1000 dilution overnight at 4C. HRP-conjugated secondary antibodies were incubated at 1:2000 for 1h. Signals were detected using Clarity Western ECL Substrate (Bio-Rad Laboratories). Band intensities were quantified using Fiji Image Studio.

#### Immunofluorescence staining

Cells were grown on #1.5H coverslips and fixed for 15 min at room temperature in 3% paraformaldehyde plus 0.2% glutaraldehyde. Samples were washed in PBS-CM (PBS containing 1 mM CaCl2 and 10 mM MgCl2), permeabilized for 5 min with 0.2% Triton X-100, and treated for 10 min with 1 mg/mL sodium borohydride to reduce aldehyde autofluorescence. Cells were blocked for 1 h in PBS-CM containing 10% goat serum and 1% BSA. Primary antibodies were diluted in antibody buffer (1% BSA, 2% goat serum, 0.05% Triton X-100, 50 mM sodium chloride and 150 mM trisodium citrate, or SSC) and incubated overnight at 4 °C. After washing in SSC containing 0.05% Triton X-100, samples were incubated for 1 h at room temperature with fluorophore-conjugated secondary antibodies, washed again, rinsed in Milli-Q water, and mounted in ProLong Diamond Antifade Mountant (Invitrogen / Thermo Fisher Scientific, Cat P36965). Coverslips were cured for 24-48 h before imaging. Antibodies and dilutions are listed in Table S1.

##### STED microscopy

2D and 3D STED images were taken using a Leica TCS SP8 STED microscope (Leica Microsystems, Wetzlar, Germany) equipped with a white-light excitation laser, 592-, 660-, and 775-nm depletion lasers, HyD detectors, and a 100×/1.40 NA oil-immersion HC PL APO CS2 objective. Time-gated detection was used for STED acquisition. Fluorophores were excited and depleted with wavelength-appropriate settings, and channels were acquired sequentially from longer to shorter emission wavelengths to minimize crosstalk. Acquisition settings were held constant within each experiment. STED stacks were deconvolved in Huygens Professional (Scientific Volume Imaging) and cropped in Fiji/ImageJ. STED images were deconvolved with Huygens Professional software (Scientific Volume Imaging, Netherlands).

### Image segmentation and classification

Following deconvolution, all channels from the 3D image stacks were isolated by manual ROI drawing in FIJI, and TIFF stacks were subjected to the adaptive, cell and channel-specific z-score filter described by Cardoen et al. (2024). For each fluorescence channel, voxel intensities below a channel-specific threshold defined as μ+zσ, where μ and σ represent the mean and standard deviation of the intensity distribution, respectively, were set to zero, using a z-score threshold of 2, to suppress low-intensity background and axial signal bleed-through associated with the anisotropic point-spread function in 3D STED. The resulting 8-bit binary masks were processed in Fiji/ImageJ using a custom Jython script and the MorphoLibJ plugin [40]. Spatially distinct structures were identified by three-dimensional connected-component labeling using 6-connectivity, whereby only voxels sharing a face or corner along the x, y, or z-axis were considered connected. Each connected component was assigned a unique integer label and stored as a 16-bit labeled object map. The resulting labeled object maps were saved as TIFF stacks for subsequent analysis.

### 3D object-based colocalization analysis

3D object measurements, colocalization analyses and graph generation were performed using custom Python scripts (generated with the aid of Google Gemini) applied to the labeled object maps generated. For each cell, spatially registered label images corresponding to DFCP1-mCherry or LC3-RFP (both henceforth named RFP for simplicity), TOMM20, SU9-Halo, and SU9-GFP channels were loaded as three-dimensional volumes. Colocalization was defined as occupancy of the same voxel coordinates by two or more masks. Object volumes and overlaps were expressed as voxel counts. Before analysis, background noise from RFP was excluded to reduce segmentation noise calculated with a two-component Gaussian Mixture Model (GMM), fitted to the log-transformed size data utilizing the Expectation-Maximization algorithm. The analytical noise threshold was subsequently established at the statistical boundary between the modeled noise and signal clusters [41]. The noise threshold defined by this model in DFCP1 objects was 84 voxels and LC3, 50 voxels. Since small mitochondria fragments were relevant for degradation studies, no noise filter was applied to the TOMM20, Halo, or GFP masks.

For Figure 3, RFP voxels colocalization relationship with mitochondrial markers Tomm20, SU9-Halo, SU9-GFP, MitoAVs (GFP+ Tomm20-) and MitoLYs (Halo+ GFP-) was calculated with the denominator as the total number of voxels belonging to retained RFP objects (overlap) in each cell expressed as percentage and the population of each mitochondrial marker positive for RFP per cell (association). For Figure 4 onwards, the RFP objects were then used as a three-dimensional volumetric spatial mask to distinguish mitochondrial regions located within RFP-positive structures from those located outside. For the Halo-reference analysis, each qualifying Halo object was divided into intra-RFP and extra-RFP regions. The intra-RFP region consisted of Halo voxels occupying the same coordinates as an RFP object. SU9/GFP and TOMM20/Halo colocalization were calculated as the number of SU9 or OMM voxels divided by the total number of Halo voxels within the respective intra-RFP region, multiplied by 100.

Docking events were defined as voxels in which an RFP object overlapped only the OMM object associated with an intact mitochondrion, while neither Halo nor GFP was present at the same voxel. Intact mitochondria were identified using SU9-Halo where at least 75% of its voxels overlapped independently with OMM and GFP simultaneously. Each spatially discontinuous connected patch was counted as an independent docking event. For each event, the script recorded the size of the docking patch, the total sizes of the corresponding RFP and OMM objects, the percentage of the OMM object occupied by the docking patch, and the percentage of the RFP object occupied by the docking patch. The number of docking events in relation to the number of all the RFP objects in a cell was then calculated as a percentage to represent a ration of RFP puncta abundance to docking phenomena.

### 2D quantification of MitoAVs and MitoLYs

To identify mitoAVs (SU9-GFP or mitoGFP devoid of Tomm20) or MitoLYs (SU9-Halo with no SU9-GFP signal), binary masks using Otsu thresholding after background subtraction by z-score filter were subtracted -Tomm20 subtracted from GFP for mitoAVs or SU9-GFP subtracted from SU9-Halo. The resulting mask was filtered for objects with >0.5 circularity and 200nm size to exclude both MDVs (mitochondria-derived vesicles) and false positives due to chromatic aberration of the mitochondrial markers. The pipeline filtered out structures smaller than 200 nm to distinguish MitoAV/LYs from MDVs based on size; MitoAV/LYs we identified are on average 0.088 µm² or ≈335 nm diameter and therefore significantly larger than the ≈160 nm limit of mitochondria-derived vesicles (MDVs) [42,43].

### Statistical analysis

Statistical analyses were performed in GraphPad Prism and Python. One-way or two-way ANOVA with Tukey’s multiple-comparison test was used as specified in the figure legends. MANOVA compared the two-dimensional distributions in Fig. 6D, and ANCOVA tested whether the relationship between ER-reporter overlap and mitochondrial overlap differed between marker groups, calculated in Python. For stacked overlap distributions, only the 0-10% and 90-100% bins were tested. Data are shown as mean ± SEM. The number of cells and independent experiments is given in each legend.

## Abbreviations

2D: two-dimensional
3D: three-dimensional
AMFR: autocrine motility factor receptor
ANCOVA: analysis of covariance
ANOVA: analysis of variance
AP: autophagosome
BAG6: BAG cochaperone 6
BFP: blue fluorescent protein
DMSO: dimethyl sulfoxide
DNM1L/DRP1: dynamin 1 like
ER: endoplasmic reticulum
ERAD: endoplasmic reticulum-associated degradation
GFP: green fluorescent protein
IVM: ivermectin
KDE: kernel density estimation
KDEL: peptide sequence K-Lysine D-Aspartic acid E-Glutamic acid L-Leucine
KO: knockout
MAP1LC3/LC3: microtubule associated protein 1 light chain 3
MANOVA: multivariate analysis of variance
mitoAV: mitochondrial autophagic vesicle
mitoGFP: mitochondria-targeted green fluorescent protein
mitoLY: mitolysosome
OMM: outer mitochondrial membrane
PBS: phosphate-buffered saline
PG: phagophore
PINK1: PTEN induced kinase 1
PRKN/Parkin: parkin RBR E3 ubiquitin protein ligase
PtdIns3K: phosphatidylinositol 3-kinase
PtdIns3P: phosphatidylinositol 3-phosphate
RFP: red fluorescent protein
RFmut: RING-finger mutant
riboMERC: ribosome-associated mitochondria-ER contact
ROI: region of interest
RRBP1: ribosome binding protein 1
SEM: standard error of the mean
siRNA: small interfering RNA
STED: stimulated emission depletion
SYNJ2BP: synaptojanin 2 binding protein
TOMM20: translocase of outer mitochondrial membrane 20
UBL4A: ubiquitin like 4A
USP13: ubiquitin specific peptidase 13
WT: wild type
ZFYVE1/DFCP1: zinc finger FYVE-type containing 1.

## References

[1] Vargas JNS, Hamasaki M, Kawabata T, et al. The mechanisms and roles of selective autophagy in mammals. Nat Rev Mol Cell Biol. 2022.

[2] Bento C, Renna M, Ghislat G, et al. Mammalian Autophagy: How Does It Work? Annual Review of Biochemistry. 2016;85.

[3] Dunn WAJ. Studies on the mechanism of autophagy: formation of the autophagic vacuole. Journal of Cell Biology. 1990;110:1923–1935.

[4] Norell PN, Campisi D, Mohan J, et al. Biogenesis of omegasomes and autophagosomes in mammalian autophagy. Biochemical Society Transactions. 2024;52(5):2145–2155.

[5] Axe EL, Walker SA, Manifava M, et al. Autophagosome formation from membrane compartments enriched in phosphatidylinositol 3-phosphate and dynamically connected to the endoplasmic reticulum. The Journal of Cell Biology. 2008;182(4):685–701.

[6] Li M, Tripathi-Giesgen I, Schulman BA, et al. In situ snapshots along a mammalian selective autophagy pathway. Proceedings of the National Academy of Sciences. 2023;120(12):e2221712120.

[7] Ridley SH, Ktistakis N, Davidson K, et al. FENS-1 and DFCP1 are FYVE domain-containing proteins with distinct functions in the endosomal and Golgi compartments. J Cell Sci. 2001;114(Pt 22):3991–4000.

[8] Hayashi-Nishino M, Fujita N, Noda T, et al. A subdomain of the endoplasmic reticulum forms a cradle for autophagosome formation. Nature Cell Biology. 2009;11(12):1433–1437.

[9] Dooley HC, Razi M, Polson HE, et al. WIPI2 links LC3 conjugation with PI3P, autophagosome formation, and pathogen clearance by recruiting Atg12-5-16L1. Mol Cell. 2014;55(2):238–52.

[10] Nakatogawa H. Mechanisms governing autophagosome biogenesis. Nature Reviews Molecular Cell Biology. 2020;21(8):439–458.

[11] Christianson JC, Jarosch E, Sommer T. Mechanisms of substrate processing during ER-associated protein degradation. Nat Rev Mol Cell Biol. 2023;24(11):777–796.

[12] Ballar P, Shen Y, Yang H, et al. The Role of a Novel p97/Valosin-containing Protein-interacting Motif of gp78 in Endoplasmic Reticulum-associated Degradation. Journal of Biological Chemistry. 2006;281(46):35359–35368.

[13] Chen B, Mariano J, Tsai YC, et al. The activity of a human endoplasmic reticulum-associated degradation E3, gp78, requires its Cue domain, RING finger, and an E2-binding site. Proceedings of the National Academy of Sciences. 2006;103(2):341–346.

[14] Das R, Mariano J, Tsai YC, et al. Allosteric Activation of E2-RING Finger-Mediated Ubiquitylation by a Structurally Defined Specific E2-Binding Region of gp78. Molecular Cell. 2009;34(6):674–685.

[15] Christianson JC, Olzmann JA, Shaler TA, et al. Defining human ERAD networks through an integrative mapping strategy. Nature Cell Biology. 2011;14(1):93–105.

[16] Fang S, Ferrone M, Yang C, et al. The tumor autocrine motility factor receptor, gp78, is a ubiquitin protein ligase implicated in degradation from the endoplasmic reticulum. Proceedings of the National Academy of Sciences. 2001;98(25):14422–14427.

[17] Liu Y, Soetandyo N, Lee J-G, et al. USP13 antagonizes gp78 to maintain functionality of a chaperone in ER-associated degradation. eLife. 2014;3.

[18] Alan P, Vandevoorde KR, Joshi B, et al. Basal Gp78-dependent mitophagy promotes mitochondrial health and limits mitochondrial ROS. Cell Mol Life Sci. 2022;79(11):565.

[19] Fu M, St-Pierre P, Shankar J, et al. Regulation of mitophagy by the Gp78 E3 ubiquitin ligase. Molecular Biology of the Cell. 2013;24(8):1153–1162.

[20] Mookherjee D, Das S, Mukherjee R, et al. RETREG1/FAM134B mediated autophagosomal degradation of AMFR/GP78 and OPA1 —a dual organellar turnover mechanism. Autophagy. 2021;17(7):1729–1752.

[21] Mukherjee R, Chakrabarti O. Ubiquitin-mediated regulation of the E3 ligase GP78 by MGRN1 in trans affects mitochondrial homeostasis. J Cell Sci. 2016;129(4):757–73.

[22] González A, Covarrubias-Pinto A, Bhaskara RM, et al. Ubiquitination regulates ER-phagy and remodelling of endoplasmic reticulum. Nature. 2023;618(7964):394–401.

[23] Zachari M, Gudmundsson SR, Li Z, et al. Selective Autophagy of Mitochondria on a Ubiquitin-Endoplasmic-Reticulum Platform. Developmental Cell. 2019;50(5):627–643.e5.

[24] Cardoen B, Vandevoorde KR, Gao G, et al. Membrane contact site detection (MCS-DETECT) reveals dual control of rough mitochondria–ER contacts. Journal of Cell Biology. 2024;223(1).

[25] Wang PT, Garcin PO, Fu M, et al. Distinct mechanisms controlling rough and smooth endoplasmic reticulum contacts with mitochondria. J Cell Sci. 2015;128(15):2759–65.

[26] Hung V, Lam SS, Udeshi ND, et al. Proteomic mapping of cytosol-facing outer mitochondrial and ER membranes in living human cells by proximity biotinylation. eLife. 2017;6.

[27] Yim WW, Yamamoto H, Mizushima N. A pulse-chasable reporter processing assay for mammalian autophagic flux with HaloTag. Elife. 2022;11.

[28] Xu Y, Liu Y, Lee JG, et al. A ubiquitin-like domain recruits an oligomeric chaperone to a retrotranslocation complex in endoplasmic reticulum-associated degradation. J Biol Chem. 2013;288(25):18068–76.

[29] Kabeya Y, Mizushima N, Ueno T, et al. LC3, a mammalian homologue of yeast Apg8p, is localized in autophagosome membranes after processing. The EMBO Journal. 2000;19(21):5720–5728.

[30] Wang Q, Liu Y, Soetandyo N, et al. A Ubiquitin Ligase-Associated Chaperone Holdase Maintains Polypeptides in Soluble States for Proteasome Degradation. Molecular Cell. 2011;42(6):758–770.

[31] de Graaf AO, van den Heuvel LP, Dijkman HB, et al. Bcl-2 prevents loss of mitochondria in CCCP-induced apoptosis. Exp Cell Res. 2004;299(2):533–40.

[32] Mlejnek P. Caspase-3 activity and carbonyl cyanide m-chlorophenylhydrazone-induced apoptosis in HL-60. Altern Lab Anim. 2001;29(3):243–9.

[33] Padman BS, Bach M, Lucarelli G, et al. The protonophore CCCP interferes with lysosomal degradation of autophagic cargo in yeast and mammalian cells. Autophagy. 2013;9(11):1862–1875.

[34] Ragimbeau R, El Kebriti L, Sebti S, et al. BAG6 promotes PINK1 signaling pathway and is essential for mitophagy. Faseb j. 2021;35(2):e21361.

[35] Tanaka A, Cleland MM, Xu S, et al. Proteasome and p97 mediate mitophagy and degradation of mitofusins induced by Parkin. J Cell Biol. 2010;191(7):1367–80.

[36] Yoshii SR, Kishi C, Ishihara N, et al. Parkin mediates proteasome-dependent protein degradation and rupture of the outer mitochondrial membrane. J Biol Chem. 2011;286(22):19630–40.

[37] Chan NC, Salazar AM, Pham AH, et al. Broad activation of the ubiquitin-proteasome system by Parkin is critical for mitophagy. Hum Mol Genet. 2011;20(9):1726–37.

[38] Wei Y, Chiang WC, Sumpter R, Jr., et al. Prohibitin 2 Is an Inner Mitochondrial Membrane Mitophagy Receptor. Cell. 2017;168(1-2):224–238 e10.

[39] Karbowski M, Youle RJ. Regulating mitochondrial outer membrane proteins by ubiquitination and proteasomal degradation. Current Opinion in Cell Biology. 2011;23(4):476–482.

[40] Legland D, Arganda-Carreras I, Andrey P. MorphoLibJ: integrated library and plugins for mathematical morphology with ImageJ. Bioinformatics. 2016;32(22):3532–3534.

[41] Dempster AP, Laird NM, Rubin DB. Maximum Likelihood from Incomplete Data Via the EM Algorithm. Journal of the Royal Statistical Society: Series B (Methodological). 1977;39(1):1–22.

[42] Sugiura A, Mclelland GL, Fon EA, et al. A new pathway for mitochondrial quality control: mitochondrial-derived vesicles. The EMBO Journal. 2014;33(19):2142–2156.

[43] Todkar K, Chikhi L, Desjardins V, et al. Selective packaging of mitochondrial proteins into extracellular vesicles prevents the release of mitochondrial DAMPs. Nature Communications. 2021;12(1).

