## Supplemental figure for "The Gp78 ubiquitin E3 ligase drives mitophagy by regulating omegasome formation"

### Supplemental figure Legend

**Supplementary Fig. 1 | IVM increases mitoAV and mitoLY area and count.** Total area, object count, and mean object size per cell are shown for mitoAVs and mitoLYs after DMSO or 10 μM IVM for 90 min. Data are mean ± SEM from $n \approx 50$ cells across 3 independent experiments. Groups were compared by non-parametic T-test. ns, not significant; *$P < 0.05$; ****$P < 0.0001$.

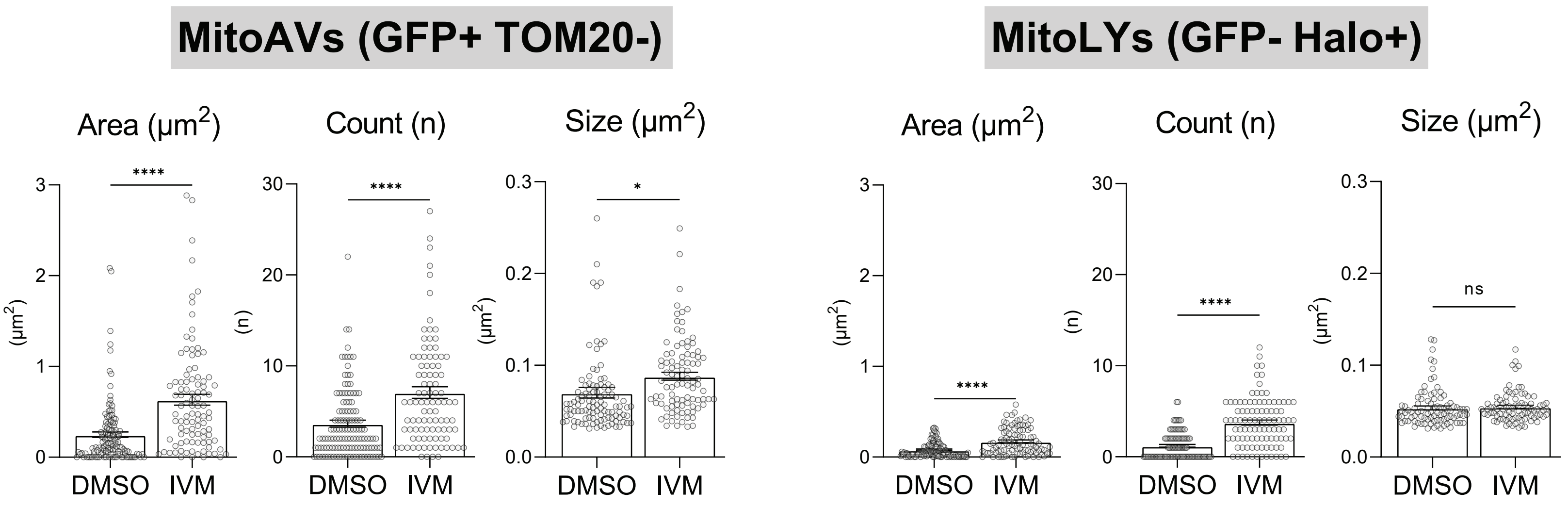


Ortiz-Silva et al Supp Figure 1
